# Effect of diesel contamination on bacterial populations in a pristine soil during rhizoremediation

**DOI:** 10.1101/2025.03.21.644530

**Authors:** Pieter van Dillewijn, Irene Hurtado-Fernández, Pablo Garrido-Peláez, Lázaro Molina, Zulema Udaondo, Félix Velando, Ana Segura

## Abstract

**Background and aims:** Contamination of soils with hydrocarbons such as diesel is of major concern as use and transport increases. In order to obtain a better understanding of how soil microorganisms face diesel during rhizoremediation of a natural soil will help us improve remediation processes.

**Methods:** We conducted microcosm experiments with a pristine soil contaminated or not with diesel and planted with clover, clover/ryegrass or non-planted controls. We measured the aromatic fraction of diesel by HPLC-DAD along time under the different experimental conditions and determined the bacterial communities in soil and the rhizosphere by amplicon sequencing of the 16S rRNA gene.

**Results:** The presence of clover accelerates the disappearance of the aromatic components of diesel but not significantly so. The bacterial diversity is lowered by diesel contamination although this decline is mitigated by the presence of plants. Bacterial community structure is mostly affected by soil type and then by diesel contamination. Actinobacteriota and Gemmatimonadota decline while Firmicutes, Bacteroidota and Proteobacteria increase with diesel contamination and in the rhizosphere. Amplified sequence variants (ASVs) belonging to *Novosphingobium* and *Sphingomonas* are especially abundant in diesel contaminated bulk soil and ASVs belonging to *Azospirillum*, *Rhizobium*, *Ohtaekwangia Faecalibacterium*, and *Subdoligranulum* are especially abundant in diesel contaminated rhizosphere.

**Conclusion:** Clover helps dissipate diesel aromatic compounds faster from a pristine soil. Clover and ryegrass mitigate the decline of bacterial diversity in soil due to diesel contamination and certain bacterial taxa which are enriched with the contaminant are identified which may play a role to remediate diesel contamination.

## Introduction

One of the most important sources of environmental contamination are from petroleum hydrocarbons and petroleum derived products such as lubricants, gasoline and diesel (Pinedo et al. 2013). In Europe alone a large percentage of contaminated sites are polluted with mineral oil, polycyclic aromatic hydrocarbons (PAHs) and volatile aromatic hydrocarbons such as benzene and toluene (BTEX) (Panagos et al. 2013; Pinedo et al. 2013). As demand and use of diesel continues to rise, contamination of soils with this petroleum derived fuel has become a major concern and as a result much effort has been directed to restore affected areas. A technology which has received increased attention to eliminate diesel and other petroleum derived contaminants from soil is rhizoremediation (Barrutia et al. 2011; Correa-Garcia et al. 2018; Hoang et al. 2021). Rhizoremediation is the remediation of pollutants by microorganisms of the rhizosphere, the soil in close contact with plant roots, which are stimulated by nutrients from plant exudates (Kuiper et al. 2004; Pilon-Smits 2005; Segura et al. 2009). The capacity to degrade hydrocarbons is widespread among bacteria but for effective rhizoremediation the characteristics of the plant are also important (Hoang et al. 2021).

Many studies have been directed at the rhizoremediation of petroleum contaminated soils (Lopez-Echartea et al. 2020; Radwan et al. 1995; Yergeau et al. 2014; Yergeau et al. 2018) but less have focused specifically at those contaminated with diesel. Diesel is made up mostly of aliphatic saturated hydrocarbons and approximately 25 % of aromatic hydrocarbons including polycyclic hydrocarbons (PAHs) listed in the EPA and EU lists of priority pollutants (Directive 2013/39/EU; Risher and Rhodes, 1995; Liang et al. 2005; Sjögren et al. 1995). To measure diesel remediation either total petroleum hydrocarbons (TPH), diesel range organics, or only the *n*-alkane or PAH constituents are determined. In a study by Palmroth et al. (2002) with a number of different plants the authors found that diesel fuel disappeared more rapidly with a legume mixture (white clover and pea). A follow-up study with scots pine or poplar and an additional application of diesel showed improved diesel removal upon the second application (Palmroth et al. 2005). High diesel removal (up to 97%) was also reported by Tesar (2002) with black poplar and herbal plants. Likewise, Kaimi et al. (2006) found enhancement of biodegradation in diesel-contaminated soil planted with ryegrass (*Lolium multiflorum* L) and Gaskin and Bentham (2010) reported the highest TPH removal with *Cymbopogon ambiguous* among the three Australian grasses tested for diesel rhizoremediation. Wojtera-Kwiczor et al. (2014) observed higher diesel removal in experiments with two rapeseed varieties and Leewis et al. (2016) found that the presence of willow (*Salix alaxensis*) removed more diesel from a contaminated soil than unplanted soil. Garrido-Sanz et al. (2019) when testing a diesel degrading consortium observed a larger decrease of TPH by inoculated alfalfa growing in diesel contaminated soil than by uninoculated plants or in unplanted soil. Lee et al. (2021; 2022) observed improved TPH removal from diesel contaminated soils planted with tall fescue. Other rhizoremediation studies with diesel contaminated soil and maize (*Zea mays*) (Chaîneau et al. 2000), maize and tall fescue (Seo and Cho, 2021) or *Megathyrsus maximus* (Uribe et al. 2022) revealed initially higher degradation rates of diesel hydrocarbons but after a longer period the biodegradation rates were the same as in unplanted soils. On the other hand, yet other diesel rhizoremediation experiments with white clover (*Trifolium repens*) and perennial ryegrass (*L. perenne)* (Barrutia et al. 2011) or *Brassica napus* (Lacalle et al. 2018) found that *n*-alkane concentrations or dissipation of TPH, respectively, were similar between planted and unplanted soils.

While generally rhizoremediation has shown positive results to remove diesel from contaminated soils it is also important to determine the effect on the microbial populations in the rhizosphere. Initially these types of studies were directed at measuring general enzyme activities, determining the number of bacteria which grow on hydrocarbons and analyzing functional or 16s rRNA gene profiles. For instance, Tesar et al. (2002) observed using T-RFLP that the diesel concentration rather than plant species caused the most marked changes in the rhizosphere microbial communities. Using Biolog MT2 assays, Palmroth et al. (2005) found higher utilization of diesel by soil bacteria in planted contaminated soils. In two different studies with Australian grasses and diesel/oil contaminated soil, an increased number of hydrocarbon using bacteria was found with all three of the plant species tested compared to uncontaminated soil (Gaskin et al. 2008; Gaskin and Bentham 2010). In the study by Barrutia et al. (2011), the authors found that contamination with diesel affected soil microbial characteristics but this effect was less in the rhizosphere of *L. perenne*. Kaimi et al. (2006) observed increased soil enzymatic activity and more bacteria in the rhizosphere of ryegrass than in unplanted diesel contaminated soil. Lacalle et al. (2018) also showed increased soil microbial activity in the presence of *B. napus*.

More recently, metagenomic or amplicon based studies have been performed to determine the bacterial communities in diesel contaminated rhizospheres. Leewis et al. (2016) found that willow had the most important effect on the bacterial community structures in contaminated soils but SIP studies indicated an important effect by fertilizer application. Seo and Cho (2021) also observed that the structure of these communities was principally affected by maize and tall fescue and by time. On the other hand, Lee et al. (2021; 2022) observed that bacterial community structures were strongly affected by the diesel concentrations as well as by time while the effect of tall fescue was less evident. Similarly, Uribe et al. (2022) also observed that time was the principal factor affecting bacterial community structures rather than the plant *M. maximus*. With regard to bacterial diversity, Khan et al. (2018) observed lower bacterial diversity in contaminated soils but higher diversity in the rhizosphere of wheat and windmill (an Australian native grass) than in bulk soil. Higher bacterial richness and diversity was also observed in the presence of tall fescue (Lee et al 2021) or followed seasonal changes (Lee et al. 2022), however, in both studies bacterial species could be correlated positively with the rhizoremediation of diesel-contaminated soil. Seo and Cho (2021) also observed higher diversity in maize and tall fescue planted diesel contaminated soil and could associate functional bacterial species with TPH degradation.

With the aim to determine which bacterial populations help plants to restore contaminated sites, we study here the changes in rhizosphere and bulk soil bacterial populations induced in a natural soil when faced with a diesel upset and how the presence of white clover (*Trifolium repens*) or together with perennial ryegrass (*Lolium perrene*) affects diesel removal from soil.

## Materials and Methods

### Chemicals

Commercial diesel was obtained from local gasoline provider. HPLC-grade solvents were supplied by Scharlau (Barcelona, Spain). Naphthalene and anthracene were obtained from Fluka (Buchs, Switzerland). Phenanthrene, biphenyl, 1-methylnaphthalene, 2-methylnaphthalene, 1,3-dimethylnaphthalene, 1,6-dimethylnaphthalene, 2,6-dimethylnaphthalene and pyrene were obtained from Sigma-Aldrich (Steinheim, Germany).

### Microcosm experiment

Microcosm experiments were set up using topsoil obtained from a natural site near Padul in Southern Spain (37.015801 N, -3.646959 E) 37°00’56.9“N 3°38’49.1“W which was sifted to 0.2 cm and mixed 1:1 (v/v) with sterile sand. The characteristics of this soil/sand mixture was the following: pH 8.57, assimilable potasium 41 p.p.m., assimilable phosphorus, 4.5 p.p.m., organic nitrogen 0.06%, organic matter content 1.37%, and CaCO_3_ content of 20.15%. The resulting mixtures were mixed with 0 (water) or 2% diesel (v/w) and allowed to acclimatize for 30 d with intermittent mixing and humidification with tap water. The microcosm experiment was initiated on June 2022 by introducing 175g soil mixture into plastic pots. Pots were planted with 30 white clover seeds (*Trifolium repens*) or together with 25 Perennial ryegrass seeds (*Lolium perenne*). Pots were introduced into mini greenhouse containers with openings and placed outside on a table covered with translucent plexiglass. Pots were watered with tap water when needed and reseeded with approx. 170 clover seeds/pot after 8 days to ensure sufficient germination. The mean daily temperature during course of the experiment was 25.16 ± 2.03 °C oscillating between 18 and 32 °C (source AEMET: Agencia Estatal de Meteorología, Spain).

Bulk soil samples were taken at initial (T=0), 28 days and 49 days and rhizosphere soil at 49 days. To obtain soil samples, plants with roots attached were removed from the pots and the remaining soil thoroughly mixed/homogenized. A portion of this bulk soil was used to determine aromatic components of diesel and another portion frozen at -20 °C for posterior DNA extraction. To obtain rhizosphere soil, roots of all the plants growing in the pots were gently shaken to remove excess bulk soil and then separated from the aerial part with sterile scissors before being introduced into a 15 ml or 50 ml tube containing 5 or 15 ml cooled sterile PBS buffer, respectively. Tubes were then vortexed for 1 min and introduced into a sonication bath (Bransonic ultrasonic cleaner) for 5 min. Roots were removed with sterile forceps and then centrifuged at 6000 rpm on a table top centrifuge for 5 minutes. Supernatant was removed and the wet pellet considered to be rhizosphere soil was frozen at -20 °C for posterior DNA extraction.

### Diesel extraction and analysis of aromatic components by HPLC-DAD

To determine and quantify the aromatic components of diesel, soil samples (3 g) were extracted with an equal volume of methanol (VWR, HPLC grade) by shaking overnight in sealed tubes followed by centrifugation for 5 minutes at 6000 rpm on a table top centrifuge. 1 ml of the methanol extract was passed through a 0.45 µm J.T. Baker H-PTFE syringe filter (VWR International LLC, PA, USA) prior to analysis by high-performance liquid chromatography (HPLC) while the remainder of the sample were air dried to determine dry weight of each soil sample. Remaining material in HPLC was performed with an Agilent 1260 Infinity II (Agilent Technologies, Santa Clara, CA, USA) equipped with a diode array detector (DAD) and a Zorbax Ecilpse PAH column (5µm, 3mm x 250 mm, Agilent Technologies). The mobile phase consisted of acetonitrile (ACN) and water MilliQ both acidified with 0.1% orthophosphoric acid. Samples were run at a flow rate of 0.85 ml/min in a gradient of 5 minutes starting from 50% (vol/vol) ACN:water to 60% ACN followed by a gradient of 20 minutes to reach 62% ACN, and then a gradient of 7 minutes to reach 100% ACN. The ACN concentration was maintained at 100% for an additional 13 minutes before dropping to 50% ACN in 1 minute to re-equilibrate at 50% ACN for an additional 15 minutes. The detector was set at 230 nm. Peak areas were used for comparison purposes and corrected for the quantity of humidity in each soil sample.

### Total DNA isolation and amplicon sequencing analysis

DNA from bulk soil and rhizosphere soil was obtained using the Fast DNA Spin Kit for Soil (MP Biomedicals LLC, Solon, Ohio, USA). Purity and DNA concentrations were determined using Nanodrop ND-1000 (Thermo Fisher Scientific) and Qubit dsDNA BR Assay kit (Live Technologies, Invitrogen, USA). Paired end amplicon sequencing of the V3-V4 region of the 16S ribosomal RNA gene with primers 5’ CCTAYGGGRBGCASCAG 3’ and, 5’ GGACTACNNGGGTATCTAAT 3’ was performed by Novogene Co, Ltd (Cambridge, United Kingdom) using Illumina technology for paired end sequencing of 250 bp. Data was provided as forward and reverse reads with barcodes and primer sequences removed.

### Bioinformatic data analysis

16S rRNA amplicon sequences were processed using QIIME2 version 2022.2 (https://qiime2.org) (Bolyen et al. 2019). Demultiplexed forward and reverse sequences were trimmed, quality filtered, denoised, joined and dereplicated using DADA2 (Callahan et al. 2016). Feature frequency of original dataset was set to at least 10 repetitions. Representative features (a.k.a ASVs or amplified sequence variants) were classified with the SILVA database v138 (Quast et al. 2013), aligned with MAFFT (Katoh and Standley 2013) and phylogenetic trees were constructed using FastTree (Price et al. 2010). Alpha diversity indices (observed features and Shannon diversity index), and beta diversity of bacterial communities were determined using QIIME2. UPGMA and principal component analysis (PCoA) based on Bray-Curtis distances, as well as PERMANOVA analysis (999 permutations) were also performed using QIIME2. UPGMA newick trees were viewed with Phylogenetic tree viewer in the ETE toolkit (http://etetoolkit.org/treeview/) (Huerta-Cepas et al. 2016).

In order to detect biomarker ASVs which showed differences between the different treatments, soil types and time points, the linear discriminant analysis (LDA) effect size (LEfSe) algorithm (Segata et al. 2011) was used in the MicrobiomeAnalyst platform (Lu et al. 2023) https://www.microbiomeanalyst.ca/MicrobiomeAnalyst/ModuleView.xhtml) with default settings for data filtering and with data normalization using cumulative sum scaling. Similarly, ANCOM (Mandal et al. 2015) was used within the QIIME2 package to determine enriched ASVs depending on conditions.

One-way analysis of variance (ANOVA) at a significance level of 0.05 were calculated using the Excel program from Microsoft Office (2019) (Microsoft).

## Results

### Elimination of diesel aromatic components in the microcosm experiment

The aliphatic saturated hydrocarbons which make up approximately 75% of diesel are mostly alkanes including *n*, *iso*, and cycloalkanes (C_10_–C_22_), and the remaining approx. 25 % are aromatic hydrocarbons including polycyclic hydrocarbons (PAHs), alkylated PAHs and alkylbenzenes (Liang et al. 2005; Sjögren et al. 1995). In this work we sought to monitor the aromatic fraction of diesel and determine how these are affected in microcosm soil experiments in the presence and absence of clover or clover with ryegrass. The aromatic components of diesel were determined by HPLC at the initial time point, after 28 days and after 49 days. According to the chromatogram obtained (See Fig. S1A) at least 25 peaks could be separated with the method used (See Materials and Methods Section). Of these, 17 of the most representative peaks were chosen for comparison purposes. Comparisons of these peaks with pure compounds for retention time and UV spectrum suggest most belong to methylated naphthalenes and possibly pyrene (Fig. S1B and C, Fig. S1 legend). Variations in the sum of the peak areas observed in the samples of the microcosm experiment are shown in Fig. 1 and of each separate peak in Fig. S2.

**Fig. 1.**
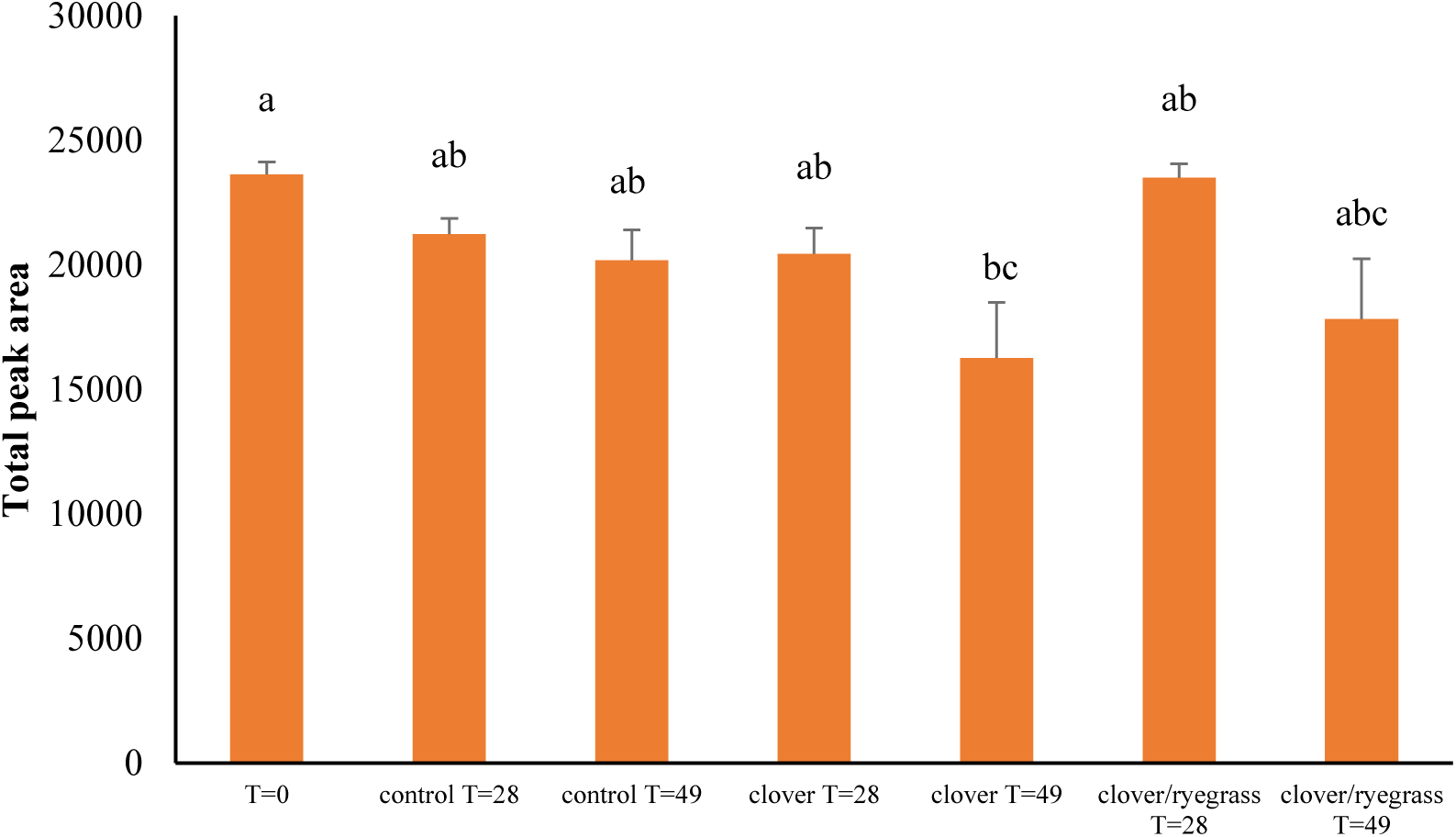
Mean total area of the 17 aromatic diesel components detected in each treatment and time point in the microcosm samples which had initially been treated with 2% diesel (v/w). Bars represent standard error of the mean and letters indicate statistical similarity according to one-way ANOVA and a post hoc Tukey’s test for pairwise comparisons (p≤0.05).

The results in Fig. 1 indicate that the strongest decrease in aromatic diesel compounds was observed in the bulk soil of pots planted with clover plants after 49 days. This decrease is statistically different to control bulk soil at T=0 but not control bulk soil after 28 or 49 days nor in pots planted with both clover and ryegrass after 49 days. The areas of each of the aromatic diesel component peaks separately (See Fig. S2), generally, share the same patterns as the total peak areas indicating that none of the compounds represented by the peaks are correlated to a specific condition or time. The presence of additional peaks forming along the time of the experiment were not detected suggesting that no new metabolites were formed from the original peaks under any of the conditions.

### Bacterial richness and diversity in diesel contaminated microcosms

To determine the response of bacterial communities towards diesel contamination in a pristine soil planted with or without clover or clover together with ryegrass, amplicon products of the V3-V4 region of the 16SrRNA gene were analyzed from gDNA extracted from three biological replicates of bulk soil and rhizosphere soil for each condition at the start (T=0) and end (T=49 days) of the experiment. After data filtering and dereplication a total of 1171787 reads associated with 6048 different amplified sequence variants (ASVs) were detected in all the replicates and conditions together.

The alpha indices indicate that in the absence of diesel the bacterial diversity, richness and evenness (Fig. 2ABC), remains similar in the control, bulk and rhizosphere soil of clover and clover/ryegrass along the time except for a slight but significant increase of bacterial richness in the rhizosphere soil of clover (Fig. 2B). On the other hand, in the presence of diesel, the bacterial diversity, richness and evenness decreases significantly after 49 days in the control, bulk and, especially, rhizosphere soil of clover. Nevertheless, this decrease is less pronounced in the bulk or rhizosphere soil of clover/ryegrass where the diversity, richness and evenness was significantly higher than in the contaminated control soil after 49 days. These results indicate that diesel has a strong negative effect on the bacterial diversity and richness in soil and that this effect of diesel contamination is mitigated in soils planted with both clover and ryegrass.

**Fig. 2.**
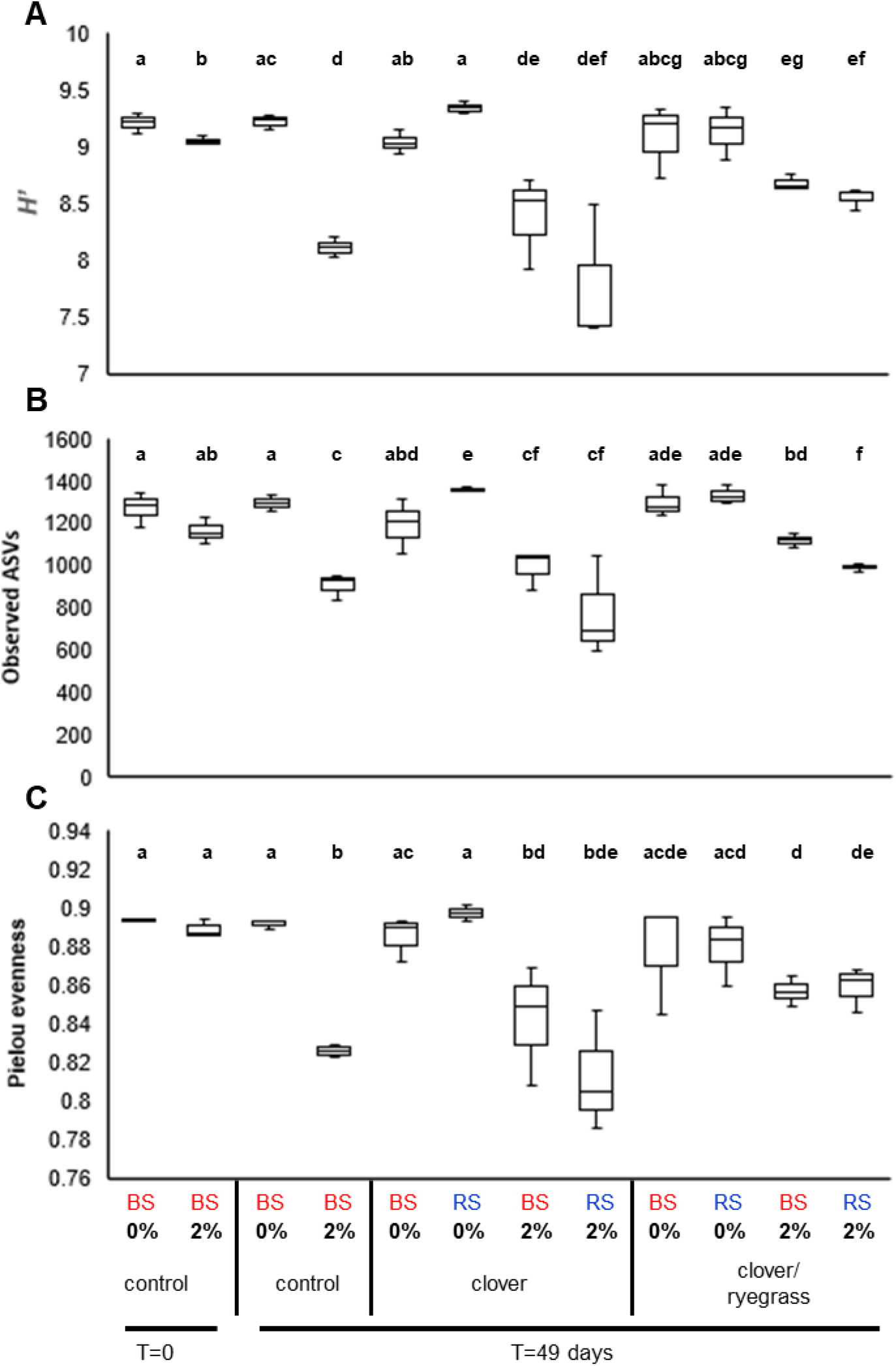
Boxplots of alpha indices for the different conditions. **a**) Shannon Index (*H*’) for bacterial diversity; **b**) Observed ASVs for bacterial richness; **c**) Pielou evenness for bacterial species homogeneity. BS: Bulk soil, RS: Rhizosphere soil; % diesel (v/w). Letters indicate significant differences according to Kruskal-Wallis (p≤0.05)

### Bacterial community structure and composition in diesel contaminated microcosms

To determine the effect of plants and diesel on the bacterial community structure, firstly the grouping of replicates was determined by UPGMA hierarchical clustering. Fig.S3 shows that most of replicates for each condition, soil type (i.e. bulk soil or rhizosphere soil), plant(s) and time grouped together except for 1 replicate of control bulk soil without diesel at 49 days and 1 replicate of clover bulk soil without diesel at 49 days which both group together with the three clover/ryegrass bulk soil samples without diesel at 49 days.

In order to determine the effects of diesel and plants on the bacterial communities, PCoA analysis (Fig. 3) was performed. The PCoA indicates that the largest effect on bacterial community structure is caused by the soil type (i.e bulk soil vs rhizosphere soil) (PERMANOVA Pseudo-F= 9.727146, p=0.001) followed by the concentration/presence of diesel (PERMANOVA Pseudo-F = 7.253863, p=0.001). The presence and type of plant also has an effect but it is less evident than the other two conditions (PERMANOVA Pseudo-F = 2.742006, p=0.002). Other factors such as time did not have an important effect (p>0.05) on the bacterial community structure.

**Fig. 3.**
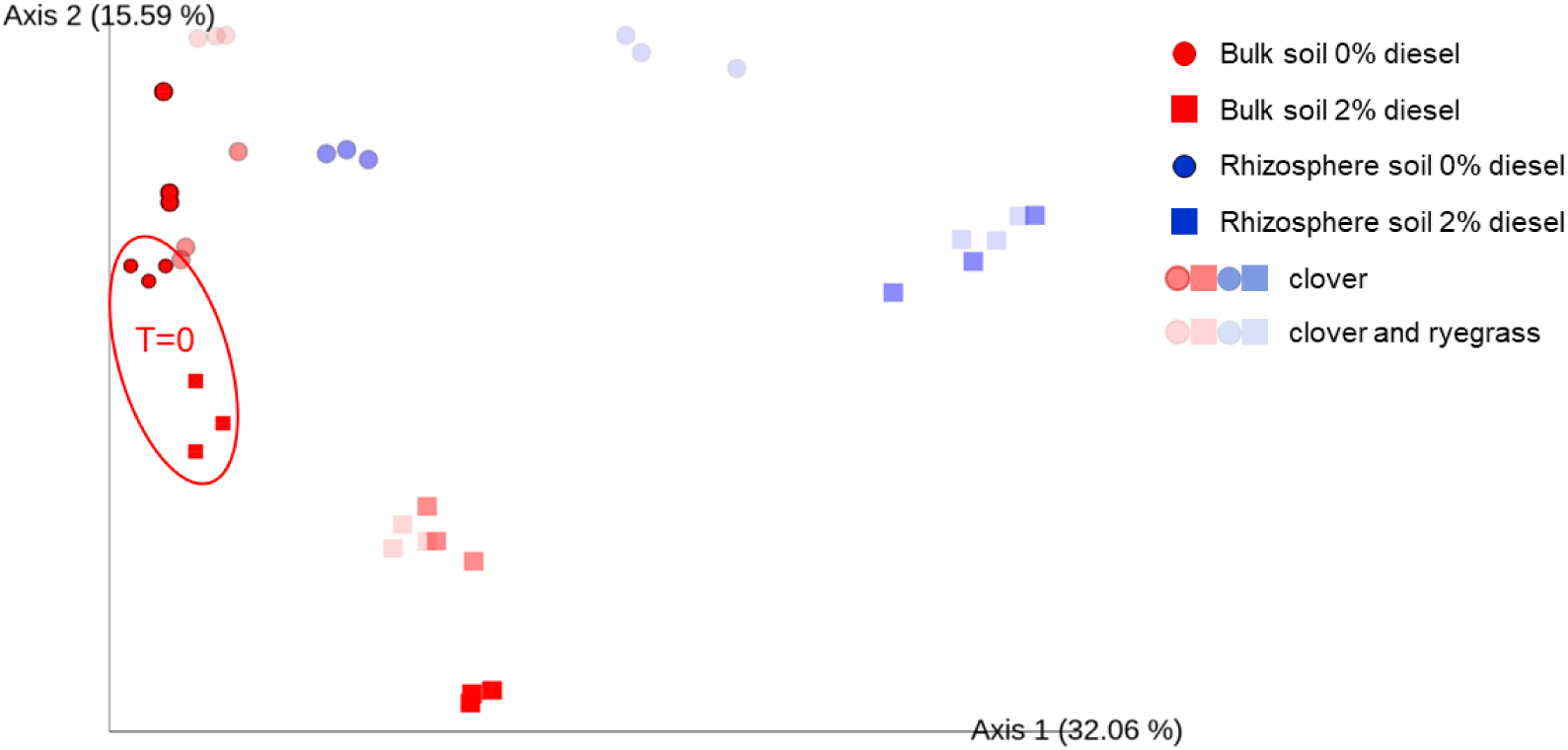
PCoA analysis based on Bray-Curtis distances. Color schemes and shading is indicated in the legend. Red symbols indicate bulk soil and blue rhizosphere soil. Spheres indicate 0% diesel and square symbols for 2% diesel. Symbols without transparency indicate control non-planted samples, 25% transparency clover samples and 50% transparency clover and ryegrass samples. Samples at T=0 are circled, all other data points are from samples taken at T=49 days.

### Impact of diesel and the soil type on the composition of taxa in the bacterial communities of the microcosm experiment

According to the taxonomic identities of the ASVs detected in the different samples at the phyla level (Fig. S4), the most abundant phylum is Actinobacteriota. The relative abundance of Actinobacteriota declined significantly (Tukey p<0.05) in the presence of 2% diesel after 49 days. In 2% diesel the relative abundance of Actinobacteriota also decreased significantly in the rhizosphere of clover/ryegrass (p<0.05) compared to bulk soil. On the other hand, the relative abundance of Firmicutes increased significantly (p<0.05) in the presence of 2% diesel and after 49 days. The relative abundance of Firmicutes also significantly increased in the rhizosphere soil of clover/ryegrass in the absence of diesel. In the case of Proteobacteria, its relative abundance increased significantly in 2% diesel and with time. As with Firmicutes, its relative abundance also increased significantly in the rhizosphere of clover/ryegrass in the absence of diesel. Gemmatimonadota decreased in the presence of 2% diesel and in the absence of diesel also decreased in the rhizosphere of clover/ryegrass. Bacteriodota, on the other hand, increased their abundance significantly in the rhizosphere of both clover and clover/ryegrass in the presence of diesel.

In order to determine differentially abundant bacterial species specific to a soil type, or to the presence of diesel or plant cover at different time points, LefSe (Segata et al. 2011), ANCOM (Mandal et al. 2015) and heatmap analysis of the most abundant ASVs were performed. Biomarker identification using Lefse (Fig. S5) indicates that the ASVs with the highest linear discriminant analysis (LDA) score in the initial bulk soil in the absence of diesel could be identified as *Nocardiodes* (Actinobacteriota) and *Sphingomonadaceae* (Proteobacteria) and in the presence of 2% diesel as *Pseudoarthrobacter* (Actinobacteriota). After 49 days in bulk soil and in the presence of diesel, ASVs identified as *Lutispora* (Firmicutes) show the highest LDA scores but also ASVs identified as *Clostridium sensu stricto* 10 (Firmicutes) as well as the proteobacterial ASVs *Novosphingobium* and *Sphingomonas*. In the bulk and rhizosphere soil of clover in the absence of diesel the highest scoring ASVs were identified as *Rhodococcus* (Actinobacteriota) and *Dongia* (Proteobacteria), respectively. However, in the presence of diesel the highest scoring ASV in clover bulk soil was identified as *Lutispora* and in clover rhizosphere *Clostridium sensu stricto* 1. In the latter condition a number of other high scoring ASVs could be identified as *Azospirillum* (Proteobacteria). In clover/ryegrass bulk or rhizosphere soil and in the absence of diesel a number of ASVs identified as *Gemmatimonadaceae* or *Rhizobium* (Proteobacteria), respectively, gave the highest LDA scores (Fig. S5). On the other hand, in the presence of diesel, ASVs identified as *Azospirillum* gave the highest LDA scores in both bulk and rhizosphere soil of clover/ryegrass. In the latter, also a number of *Faecalibacterium* and *Subdoligranulum* ASVs (both Firmicutes) gave high LDA scores. Many of the same ASVs found as possible biomarkers also appeared in the ANCOM analysis as differentially abundant (Table S1) especially the actinobacterial ASVs *Nocardiodes* which were initially abundant but no longer so after 49 days and the large number of Firmicutes ASVs identified as *Faecalibacterium* and *Subdoligranulum* which were more abundant in rhizosphere soil than in bulk soil. Noteworthy too, are the number of Bacteroidota ASVs which were more abundant in uncontaminated soil and in the rhizosphere especially an ASV identified as *Ohtaekwangia koreensis* which was found differentially more abundant both in the rhizosphere and in planted soils and the proteobacterial ASV identified as *Rhizobium* in planted soil. The enrichment of a proteobacterial ASV identified as *Novosphingobium*, the actinobacterial genus *Cellumonas* and the Patescibacteria *TM7a* was detected by ANCOM to be enriched in diesel contaminated soils (Table S1). Most of these ASVs can be corroborated in the heatmap to be especially abundant (Fig. 4) which includes the ASVs identified by LefSe and ANCOM as well as other ASVs which are highly abundant in the microcosm assay. Altogether, the results in Fig. 4 indicate the especially high abundance of actinobacterial ASVs (mostly *Nocardiodes*) and an proteobacterial ASV *Sphingomonadaceae* in the initial control samples and ASVs belonging to *Rubrobacter, Gemmatimonadaceae* and *Chitinophagaceae* in uncontaminated bulk soil at the end of the experiment. A set of Firmicutes ASVs, *Lutispora* and *Clostridium sensu stricto* 10, as well as proteobacterial ASVs *Novosphingobium*, and *Sphingomonas* show a predilection for contaminated bulk soil. Also noteworthy are a number of proteobacterial ASVs, *Azospirillum* and *Rhizobium*, Firmicutes ASVs identified as *Faecalibacterium* and *Subdoligranulum*, and the Bacteroidota ASV, *Ohtaekwangia koreensis*, which are particularly abundant in contaminated rhizosphere soils (Fig. 4).

**Fig. 4.**
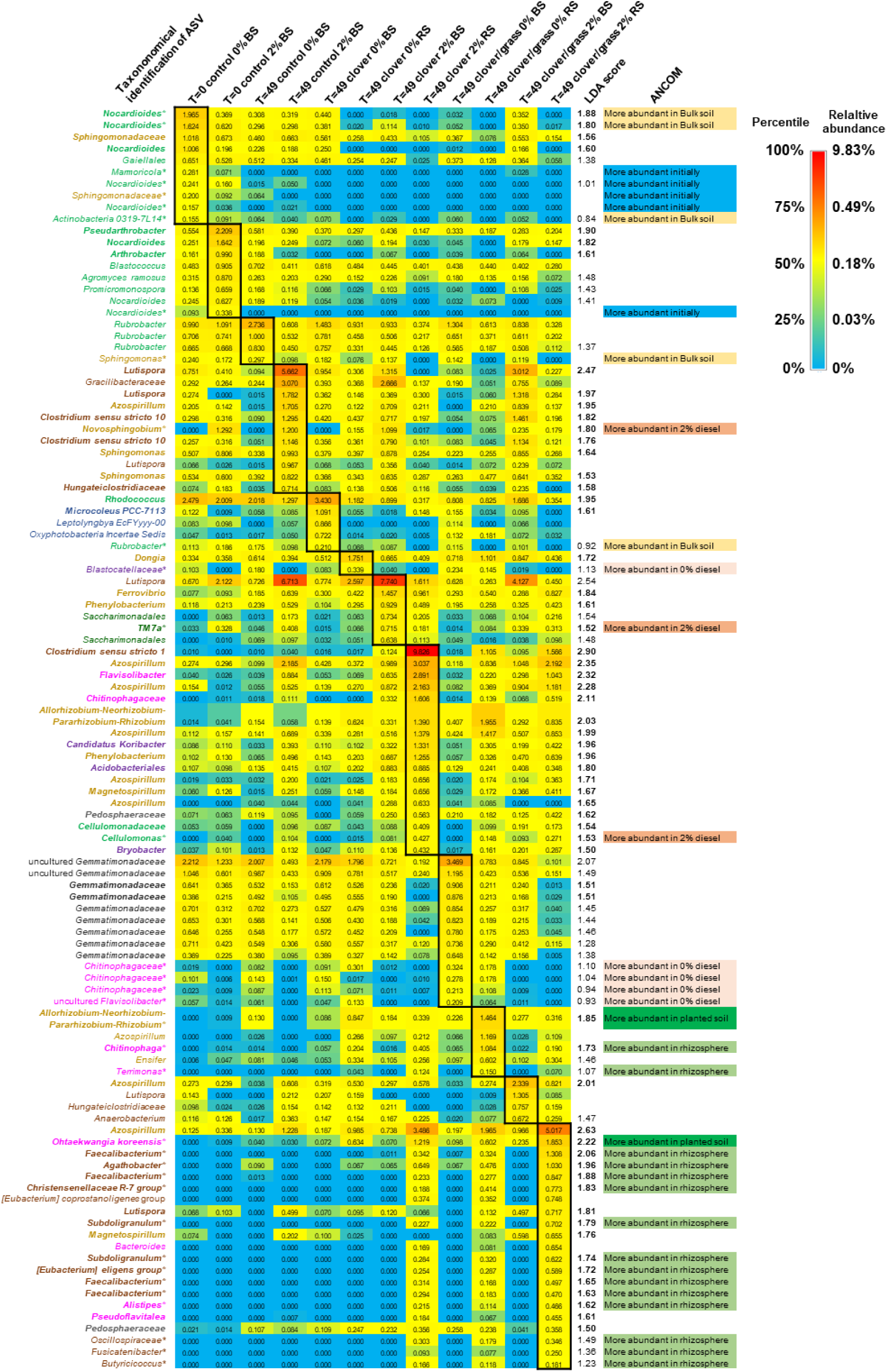
Heatmap of most frequent ASVs (> 0.6%) and/or those described in the LefSe (in bold) analysis of Fig. S5 and/or ANCOM analysis (*) of Table S1. LDA scores were calculated and ANCOM results are also indicated. Those ASVs most abundant for each sample condition are boxed. BS bulk soil, RS rhizosphere soil. Letter colors of ASV taxa names correspond to their identification at the phyla level using the same color scheme as in the legend of Fig. S4.

## Discussion

The rhizoremediation experiments with a diesel contaminated natural soil performed in this work with white clover (*T. repens*) or together with ryegrass (*L. perenne*) showed that the strongest removal of the aromatic fraction of diesel was found in the soil associated with clover after 49 days followed by that detected in soil planted with clover with ryegrass. However, in neither case was the observed decrease significantly different from that observed with unplanted soil after the same period of time. In a study by Palmroth et al. (2002) in which they compared diesel removal by a legume mixture (including *T. repens*), a grass mixture (including *L. perenne*), and other plants, the authors observed the fastest disappearance by the legume mixture but removal by the grass mixture was similar to that observed in unplanted soil. Similar results were found by Barrutia et al. (2011) with clover or ryegrass and diesel contaminated soil as after 5 months the amount of total *n*-alkanes in soil planted with clover was less, but not significantly so, than in soil planted with ryegrass or unplanted soil. Therefore, our results with respect to diesel dissipation are similar to previous studies using the same plant species.

Palmroth et al. (2002) also compared plant growth of the legume and grass mixtures and observed that the dry weight of legumes was 67% of that of plants grown under control conditions without diesel while grass grew even less with 47% of the dry weight under control conditions. In the study by Barrutia et al. (2011), they found that clover growth was severely affected in the presence of diesel but not that of ryegrass. Our results are largely in line with these observations as the presence of diesel affected the growth of both plants but especially that of clover (Fig. S6). Some plants can take up petroleum hydrocarbons although this capacity and the amounts depend on the plant species (Hunt et al. 2019). The only study of diesel uptake by white clover and ryegrass was that performed by Palmroth et al. (2002). In that study the legume mix did not take up detectable amounts of diesel while grass appeared to take up a little which led the authors to conclude that phytoaccumulation is not a significant diesel removal strategy. This indicates that the diesel dissipation observed in our experiments apart from evaporation of the more volatile elements of diesel is due to rhizoremediation by the microbial activity stimulated by the presence of the plant.

In order to determine to what extent the microbial and more specifically the bacterial community is involved in diesel dissipation, the presence and the effect of diesel on these populations was investigated. Our results indicate that the presence of diesel lowers bacterial richness and diversity in unplanted soil as well as in the bulk and rhizosphere soil of clover and clover with ryegrass. There is a tendency, compared to unplanted soil, for the bulk soil of clover-planted contaminated soil to show slightly higher values for these indexes, while the rhizosphere shows lower values. In contrast, in soil planted with both clover and ryegrass, both compartments (i.e., bulk soil and rhizosphere) show significantly higher values. This indicates that the presence of both plants can mitigate the negative effect of diesel on soil bacterial populations. When compared to other studies with diesel or petroleum, generally, the presence of petroleum hydrocarbons had a negative effect on bacterial richness and diversity. In the presence of diesel, the studies by Khan et al. (2018), Uribe et al. (2022), and Mitter et al. (2021) observed lowered bacterial diversity and richness compared to uncontaminated soil. Similarly, the presence of petroleum lowered bacterial diversity and richness (Eze et al. 2021; Liu et al. 2024; Mafiana et al. 2021) but not always (Shen et al. 2022; Tardif et al. 2016). On the other hand, under contaminated conditions the presence or absence of plants have been reported to give different effects. For instance, with diesel Khan et al. (2018) observed higher diversity in the rhizosphere of wheat and windmill and Lee et al. (2021) higher richness and diversity with the presence of tall fescue. Seo and Cho (2021) also observed higher diversity and richness in the rhizosphere of maize and tall fescue in diesel contaminated soil, but Uribe et al. (2022) observed less in the diesel contaminated rhizosphere of *Megathyrsus maximus*. Similarly, with petroleum and the presence of plants, Lopez-Echartea et al. (2020) observed that along three years that as the TPH values decreased both bacterial richness and diversity increased in the rhizosphere of poplar. Tardif et al. (2016) found that bacterial diversity was higher in the rhizosphere than in bulk petroleum contaminated soil but Bell et al. (2014) observed lowered bacterial diversity in willow rhizosphere of highly petroleum contaminated soil and no effect in less or non-contaminated soil. Hou et al. (2015) did not observe an effect of the presence of the plant (*Festuca arundinaceae*) on bacterial richness or diversity in petroleum contaminated soil. Altogether these studies indicate that while diesel has generally a negative effect on bacterial populations in the soil, the presence of the plant can either mitigate this effect, have no effect or exasperate this effect. In the present study the presence of clover and ryegrass mitigated the negative effect of diesel. The decrease in diversity by diesel contamination could be an indication of microbial composition turnover, allowing more specialized taxa to thrive (Jesus et al. 2021). In those cases, where the presence of the plant mitigates this effect, possibly the plant provides conditions which protect the microbial communities from the toxicity of diesel and/or provide alternative carbon sources to maintain non specialists.

When looking at the effect of diesel, the presence of the plants and the soil compartment on the bacterial community structure, our results indicate that the soil type, i.e. bulk soil or rhizosphere soil, is the major determinant followed by the presence of diesel. The presence or type of plant had a less important effect while time had no discernable effect. Compared to other studies, the diesel concentration and time had stronger effects than plants in the studies by Lee et al. (2021, 2022), whereas in the studies by Leewis et al. (2016) and Seo and Cho (2021), the effects were vice versa, or the impact was mixed, with time being the most important factor followed by the presence of the plant (Uribe et al. 2022).

In order to determine which members of the bacterial community from the natural soil could be affected by diesel or the presence of the plant, we found that the abundance of members belonging to the phyla Actinobacteriota and Gemmatimonadota declined while Firmicutes and Bacteriodota increased in the presence of 2% diesel and in the rhizosphere. Proteobacteria also increased in the presence of 2% diesel but in the rhizosphere the abundance of this phylum increased more when diesel was absent. These tendencies are reflected by the most abundant amplified sequence variants (ASVs) in which the actinobacterial *Nocardiodes* declined in time while actinobacterial *Rubrobacter*, the Gemmatinonadota *Gemmatimonadaceae* and Bacteroidota *Chitinophagaceae* declined in diesel contaminated rhizosphere while increasing in uncontaminated bulk soil. On the other hand, several ASVs belonging to Firmicutes, Proteobacteria and Bacteroidota increased with diesel contamination. A closer look at these ASVs could indicate specialist bacterial species which thrived because they could tolerate or degrade diesel compounds especially in the rhizosphere to aid the rhizoremediation process.

Among the Firmicutes ASVs which showed increased abundance in contaminated bulk soil included several belonging to *Lutispora* and *Clostridium sensu stricto* 10. A *Lutispora* ASV was correlated to benzo(a)pyrene removal in a contaminated paddy soil (Li and Chang, 2022), but generally this genus has not been associated with diesel or PAH contamination. *Clostridium* have been reported to form part of a consortium in an enrichment culture performed with diesel and oil spill contaminated sediment (Rodriguez et al. 2023) however how these obligate anaerobes interact with the fuel remains unknown. Several Firmicutes ASVs were also detected to be enriched in contaminated rhizosphere soil. These included several ASVs belonging to *Faecalibacterium* and *Subdoligranulum* which are better known as important members of the gut microbiome and as strict anaerobes. However, *Subdoligranulum* have been found in an agricultural soil (Wongkiew et al. 2022) and in soil from a natural park (Velez-Martinez et al. 2024) while *Faecalibacterium* have been found in the soil of a subtropical evergreen broad-leaved forest (Li et al. 2020). Nevertheless, neither taxa have been associated with hydrocarbon contaminated soil or to hydrocarbon degradation previously. Possibly, in the microcosm experiments, the rhizosphere soil was more anoxic than bulk soil and these taxa are more resistant to the contaminant allowing them to proliferate using plant exudated nutrients rather than by degrading diesel.

In the present work, the proteobacterial ASVs *Novosphingobium* and *Sphingomonas* showed increased abundance in diesel contaminated bulk soil. Many *Novosphingobium* strains have been described in literature which degrade PAHs, for instance, *Sphingomonas*(*Novosphingobium*) *aromaticivorans* (Shi et al. 2001), *N. pentaromativorans* (Lyu et al. 2014), *Novosphingobium panipatense* (Chettri and Singh 2019), *Novosphingobium* sp. HR1a (Segura et al 2017, 2021), and *Novosphingobium* sp HS2a (Rodriguez-Conde et al. 2016) which degrade a wide range of PAHs such as naphthalene, anthracene, phenanthrene, pyrene, chrysene, fluoranthene, acenaphthene, and methylnaphthalene. Genomic, metagenomic and metatranscriptomic studies have revealed a number of *Novosphingobium* enzymes for aromatic hydrocarbon degradation (Garrido-Sanz et al. 2019; Lyu et al. 2014; Pandolfo et al. 2024; Segura et al. 2017; Segura et al. 2021) as well as for aliphatic hydrocarbon degradation (Garrido-Sanz et al. 2019; Pandolfo et al. 2024). With respect to rhizoremediation, *Novosphingobium* HR1a was shown to proliferate better on clover roots than on those from ryegrass or grass and to eliminate a greater amount of phenanthrene in microcosms planted with clover than in those with the other plants (Molina et al. 2021). Recently, bioaugmentation of tall fescue (*Festuca arundinacea*) with *Novosphingobium* sp CuT1 in soil contaminated artificially with copper, lead and diesel showed a positive correlation with diesel removal (Lee et al. 2023). Therefore, it is possible that local *Novosphingobium* strains in the natural soil assayed in our study also play a role in the rhizoremediation of diesel hydrocarbons observed.

Numerous *Sphingomonas* strains have been isolated from hydrocarbon contaminated soils and rocks which degrade both aliphatic and aromatic hydrocarbon contaminants (Aislabie et al. 2000; Alonso-Gutierrez et al. 2009; Baraniecki et al. 2002; Semenova et al. 2023; van Herwijnen et al. 2003; Zhang et al. 2014). In a study with a contaminated soil acclimatized with phenanthrene, Gu et al. (2023) observed that among the enriched genera included *Sphingomonas*. Garrido-Sanz et al. (2019) observed by metagenomics of a diesel degrading consortium that naphthalene 1,2-dioxygenases as well as alkane degradation genes such as *alkB* (alkane 1-monoxygenase) and *ladA* (long chain alkane monoxygenase) were associated to *Sphingomonas*. *Sphingomonadales* were the major taxa performing the first steps of phenanthrene degradation in phenanthrene contaminated soil planted or not with Italian ryegrass (*Lolium multiflorum*), suggesting their critical role to initiate *in situ* PAH remediation (Thomas et al. 2019). Increased expression of *Sphingomonas* specific naphthalene dioxygenase and aromatic ring hydroxylating dioxygenase genes was observed in the rhizosphere of *Salix purpurea* growing in petroleum hydrocarbon and PCB contaminated soil (Page et al. 2015). Mukherjee et al. (2015) described the temporal shifts of the taxonomic and functional bacterial population in unplanted and planted (with poplar) after an oil truck leak. They found that *Sphingomonas* type extradiol dioxygenases highly dominated early-phase communities. Similarly, *Sphingomonas* was found to be one of the most dominant genera during the rhizoremediation of diesel contaminated soil by tall fescue (Lee et al. 2022). Altogether, these studies indicate that the abundant *Sphingomonas* ASVs are likely candidates for diesel degradation in the bulk soil of our experiments.

Among the ASVs which were enriched in contaminated rhizosphere, besides the Firmicutes ASVs mentioned above, include *Azospirillum*, *Rhizobium* and *Ohtaekwangia*. The use of *Azospirillum* for bioremediation has been reviewed by Cruz-Hernandez et al. (2022) and are especially interesting for their plant growth promoting capacity to fix nitrogen. *Azospirillum* have been found to be abundant in petroleum polluted soils (García-García et al. 2023) and Eckford et al. (2002) describe an *A. brasiliense* strain isolated from fuel contaminated soil that fixes nitrogen but does not grow with jet fuel or hydrocarbons. Wu et al. (2021) described two *Azospirillum* isolates which fix nitrogen and remove 36% heavy oil as well as producing biosurfactants. Also, inoculation of *Festuca* or *Dactylis* with *Azospirillum* and *Pseudomonas* resulted in positive effects for PAH elimination from PAH and diesel contaminated soils (Galazka and Galazka 2015; Galazka et al. 2012). Therefore, the enriched *Azospirillum* ASVs observed in our experiments may possibly be playing a more important role in aiding plant growth in the contaminated rhizosphere rather than for hydrocarbon degradation. A similar role may also be associated to *Rhizobium* ASVs which increased their abundance in contaminated rhizosphere soil where they may participate as the nitrogen fixing symbionts in clover nodules. Nevertheless, *Rhizobium* and *Sinorhizobium* isolates have been described which degrade phenanthrene (Bodour et al. 2003; Muratova et al. 2014) and even *Rhizobium* endophytes which could grow on kerosene, octanol, and motor oil and mineralize toluene and naphthalene (Lumactud et al. 2016). Lee et al. (2021) found genera including *Rhizobium* to be positively linked to diesel-contaminated soil rhizoremediation with tall fescue. *Allorhizobium-Neorhizobium-Rhizobium* formed part of a consortium aimed at degrading total petroleum hydrocarbons based on an enrichment culture using diesel as sole carbon and energy source (Pandolfo et al. 2024). Therefore, as well as playing a beneficial role for plant growth especially in the case of clover, the *Rhizobium* ASVs in our experiments could also possibly be involved in diesel degradation.

Bacteroidetal *Ohtaekwangia* have often been found associated with plants (Gu et al. 2017) including as root endophytes of ryegrass (Gaggia et al. 2013). *Ohtaekwangia* was one of the enriched genera which were significantly more abundant in the rhizosphere of *M. maximus* which had been used to remove petroleum hydrocarbons from a diesel contaminated soil (Uribe et al. 2022). Amongst the genera which increased their relative abundance in ryegrass planted soil contaminated with cadmium and pyrene also included *Ohtaekwangia* (Li et al. 2021). *Ohtaekwangia* was found to be positively correlated with the removal of C21–C34 petroleum hydrocarbon fractions by tall fescue grown in aged petroleum contaminated soil (Hou et al. 2015). As *Ohtaekgwangia* genes have not been found to be associated with hydrocarbon degradation, it remains unknown what role it plays in the contaminated rhizosphere but it is clear that its enrichment under these conditions in not unusual.

The present study of the microbiome of soil and rhizosphere in contaminated natural soil suggests that several tolerant and possibly biodegradative taxa become enriched even when the soil had no previous history of hydrocarbon contamination. As plant cover was shown to accelerate dissipation of the aromatic components of diesel and the plants used hardly accumulate diesel components by themselves, the observed accelerated dissipation is likely due to rhizoremediation by the enriched taxa. This adds new knowledge of how natural microbial populations react and aid remediation by plants and therefore provides new ways which could be used to improve the removal of diesel and restore contaminated sites for instance by manipulating microbial communities (Thijs et al. 2017) in the rhizosphere to enhance bioremediation rates by using a consortium of the identified enriched taxa.

## Acknowledgements

This study was funded by grant PID2020-116766GB-I00 funded by MCIN/AEI/ 10.13039/501100011033 and by “ERDF A way of making Europe” and by grant TED2021-129398B-I00 funded by MCIN/AEI/10.13039/501100011033/ and by the “European Union NextGenerationEU/PRTR”.

