## Supplementary Material for "Effect of diesel contamination on bacterial populations in a pristine soil during rhizoremediation"

**Table S1** Features or ASVs differentially abundant under different conditions according to ANCOM.

| | | | Percentile | | | | | | | | | |
|---|---|---|---|---|---|---|---|---|---|---|---|---|
| Phylum | Taxon | W value | 0% | 50% | 100% | 0% | 50% | 100% | 0% | 50% | 100% | Characteristic |
| **Features different for concentration** | | | 0% diesel | 0% diesel | 0% diesel | 2% diesel | 2% diesel | 2% diesel | | | | |
| *Bacteroidota* | *Chitinophagaceae* | 5608 | 1 | 65 | 128 | 1 | 1 | 12 | | | | More abundant in 0% |
| *Acidobacteriota* | *Blastocatellaceae* | 5604 | 1 | 51 | 130 | 1 | 1 | 34 | | | | More abundant in 0% |
| *Bacteroidota* | *Chitinophagaceae* | 5500 | 1 | 40 | 117 | 1 | 1 | 12 | | | | More abundant in 0% |
| *Bacteroidota* | uncultured *Flavisolibacter* | 5459 | 1 | 27 | 67 | 1 | 1 | 15 | | | | More abundant in 0% |
| *Bacteroidota* | *Chitinophagaceae* | 5141 | 1 | 25 | 83 | 1 | 1 | 11 | | | | More abundant in 0% |
| *Patescibacteria* | *TM7a* | 5275 | 1 | 14 | 42 | 21 | 118 | 345 | | | | More abundant in 2% diesel |
| *Actinobacteriota* | *Cellulomonas* | 5244 | 1 | 1 | 76 | 1 | 34 | 140 | | | | More abundant in 2% diesel |
| *Proteobacteria* | *Novosphingobium* | 5182 | 1 | 1 | 57 | 1 | 197 | 479 | | | | More abundant in 2% diesel |
| **Features different for plant** | | | clover | clover | clover | clover/grass | clover/ ryegrass | clover/ ryegrass | no plant | no plant | no plant | |
| *Proteobacteria* | *Allorhizobium-Neorhizobium-Pararhizobium-Rhizobium* | 5647 | 21 | 90 | 498 | 37 | 105 | 754 | 1 | 1 | 50 | More abundant in planted soil |
| *Bacteroidota* | *Ohtaekwangia koreensis** | 5086 | 9 | 63 | 497 | 6 | 155 | 1040 | 1 | 1 | 24 | More abundant in planted soil |
| **Features different for soil type** | | | Bulk Soil | Bulk Soil | Bulk Soil | Rhizosphere Soil | Rhizosphere Soil | Rhizosphere Soil | | | | |
| *Actinobacteriota* | *Nocardioides* | 5902 | 1 | 101 | 832 | 1 | 1 | 1 | | | | More abundant in Bulk soil |
| *Actinobacteriota* | *Nocardioides* | 5866 | 11 | 106 | 631 | 1 | 1 | 21 | | | | More abundant in Bulk soil |
| *Actinobacteriota* | *Rubrobacter* | 5608 | 19 | 37 | 87 | 1 | 1 | 25 | | | | More abundant in Bulk soil |
| *Actinobacteriota* | *Actinobacteria 0319-7L14* | 5538 | 1 | 18 | 73 | 1 | 1 | 1 | | | | More abundant in Bulk soil |
| *Proteobacteria* | *Sphingomonas* | 5335 | 1 | 50 | 168 | 1 | 1 | 51 | | | | More abundant in Bulk soil |
| *Firmicutes* | *[Eubacterium] eligens* group | 5815 | 1 | 1 | 1 | 1 | 88 | 300 | | | | More abundant in rhizosphere |
| *Firmicutes* | *Subdoligranulum* | 5797 | 1 | 1 | 1 | 1 | 76 | 328 | | | | More abundant in rhizosphere |
| *Firmicutes* | *Subdoligranulum* | 5772 | 1 | 1 | 1 | 1 | 109 | 319 | | | | More abundant in rhizosphere |
| *Firmicutes* | *Faecalibacterium* | 5740 | 1 | 1 | 1 | 1 | 61 | 235 | | | | More abundant in rhizosphere |
| *Bacteroidota* | *Terrimonas* | 5734 | 1 | 1 | 1 | 1 | 22 | 126 | | | | More abundant in rhizosphere |
| *Firmicutes* | *Christensenellaceae R-7* group | 5710 | 1 | 1 | 1 | 1 | 98 | 401 | | | | More abundant in rhizosphere |
| *Firmicutes* | *Oscillospiraceae UCG-002* | 5702 | 1 | 1 | 1 | 1 | 84 | 176 | | | | More abundant in rhizosphere |
| *Firmicutes* | *Faecalibacterium* | 5658 | 1 | 1 | 13 | 1 | 72 | 535 | | | | More abundant in rhizosphere |
| *Firmicutes* | *Faecalibacterium* | 5639 | 1 | 1 | 1 | 1 | 68 | 262 | | | | More abundant in rhizosphere |
| *Bacteroidota* | *Chitinophaga* | 5624 | 1 | 7 | 29 | 26 | 104 | 523 | | | | More abundant in rhizosphere |
| *Bacteroidota* | *Alistipes* | 5617 | 1 | 1 | 1 | 1 | 48 | 216 | | | | More abundant in rhizosphere |
| *Bacteroidota* | *Ohtaekwangia koreensis** | 5602 | 1 | 15 | 134 | 94 | 285 | 1040 | | | | More abundant in rhizosphere |
| *Firmicutes* | *Faecalibacterium* | 5587 | 1 | 1 | 11 | 1 | 90 | 657 | | | | More abundant in rhizosphere |
| *Firmicutes* | *Butyricicoccus* | 5439 | 1 | 1 | 1 | 1 | 43 | 110 | | | | More abundant in rhizosphere |
| *Firmicutes* | *Agathobacter* | 5388 | 1 | 1 | 83 | 1 | 155 | 647 | | | | More abundant in rhizosphere |
| *Firmicutes* | *Fusicatenibacter* | 5316 | 1 | 1 | 1 | 1 | 26 | 132 | | | | More abundant in rhizosphere |
| **Features different for time** | | | 0 days | 0 days | 0 days | 49 days | 49 days | 49 days | | | | |
| *Actinobacteriota* | *Nocardioides* | 5716 | 44 | 53 | 102 | 1 | 1 | 25 | | | | More abundant initially |
| *Proteobacteria* | *Sphingomonadaceae* | 5605 | 27 | 40 | 93 | 1 | 1 | 30 | | | | More abundant initially |
| *Actinobacteriota* | *Nocardioides* | 5590 | 1 | 65 | 119 | 1 | 1 | 1 | | | | More abundant initially |
| *Actinobacteriota* | *Nocardioides* | 5355 | 10 | 28 | 62 | 1 | 1 | 22 | | | | More abundant initially |
| *Actinobacteriota* | *Marmoricola* | 5255 | 1 | 51 | 110 | 1 | 1 | 29 | | | | More abundant initially |
| *Actinobacteriota* | *Nocardioides* | 5115 | 25 | 29 | 39 | 1 | 1 | 21 | | | | More abundant initially |

* This specific ASV is identical in the two comparisons

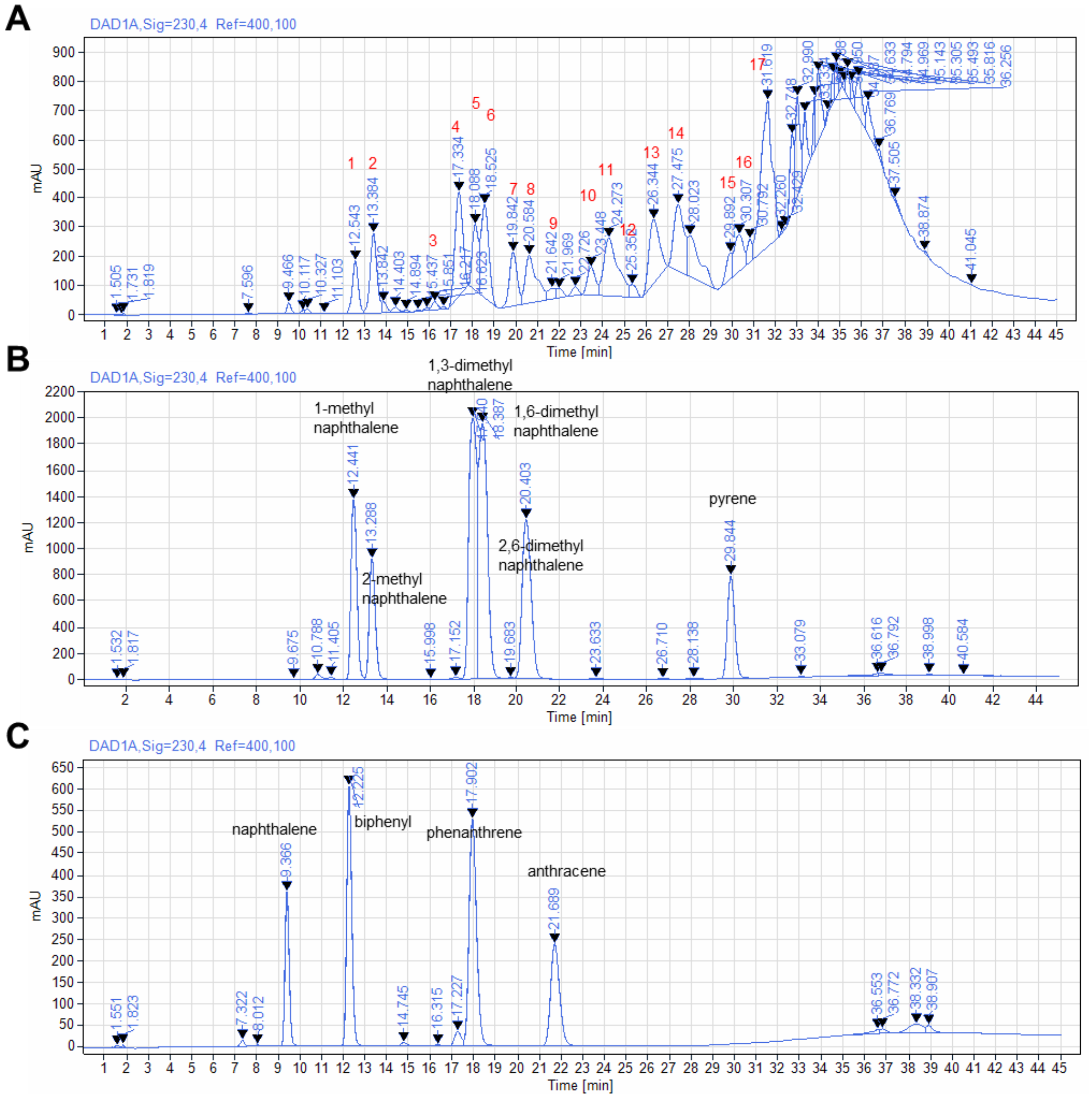


**Fig. S1** HPLC chromatograms of A) aromatic diesel components of 1% diesel diluted into methanol (Numbers indicate the peaks chosen for further analyses); B) pure methylated naphthalenes and pyrene (each 100 ppm); and C) pure naphthalene, phenanthrene, anthracene and biphenyl (each 100 ppm). Similar retention times and UV spectra indicate similarities of peak 1 with 1-methylnaphthalene, peak 2 with 2-methylnaphthalene, peak 5 with 1,3-dimethylnaphthalene, peak 6 with 1,6-dimethyl naphthalene, peak 7 with 2,6-dimethylnaphthalene and peak 15 with pyrene.

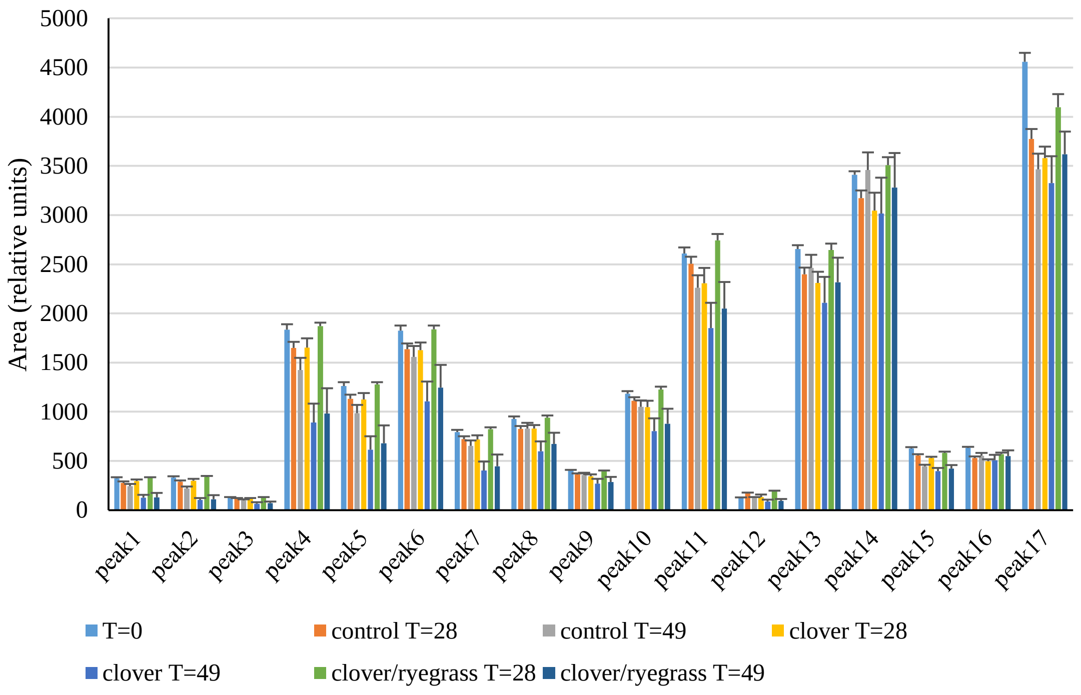


**Fig. S2** Mean area of each of the 17 peaks of aromatic diesel components found in each type of sample. Bars represent standard error of the mean.

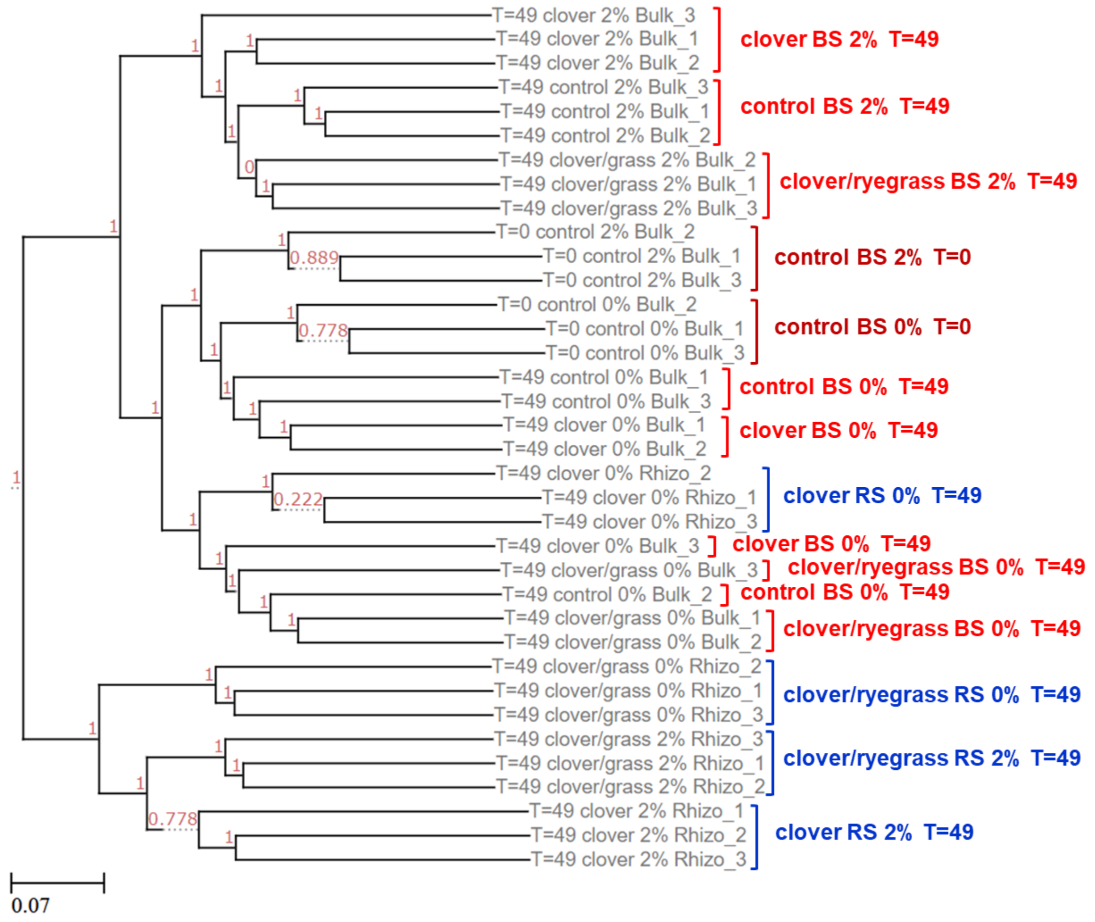


**Fig. S3** Hierarchical clustering with UPGMA based on Bray-Curtis distances. BS is bulk soil, RS is rhizosphere soil. % diesel is indicated. T is time in days.

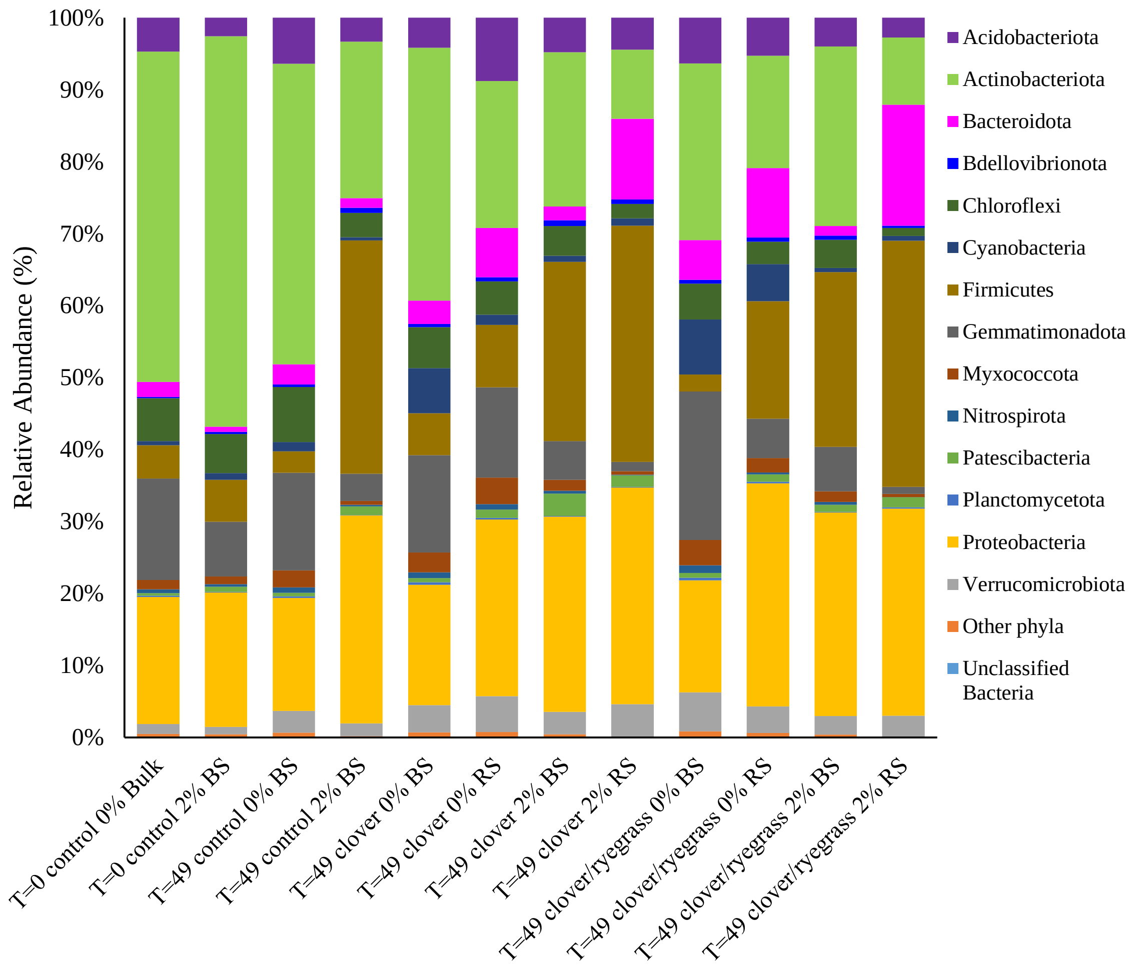


**Fig. S4** Mean relative abundances of the different bacterial phyla (>0.5% abundance) in the three replicate samples for each condition and time. BS indicates bulk soil and RS rhizosphere soil.

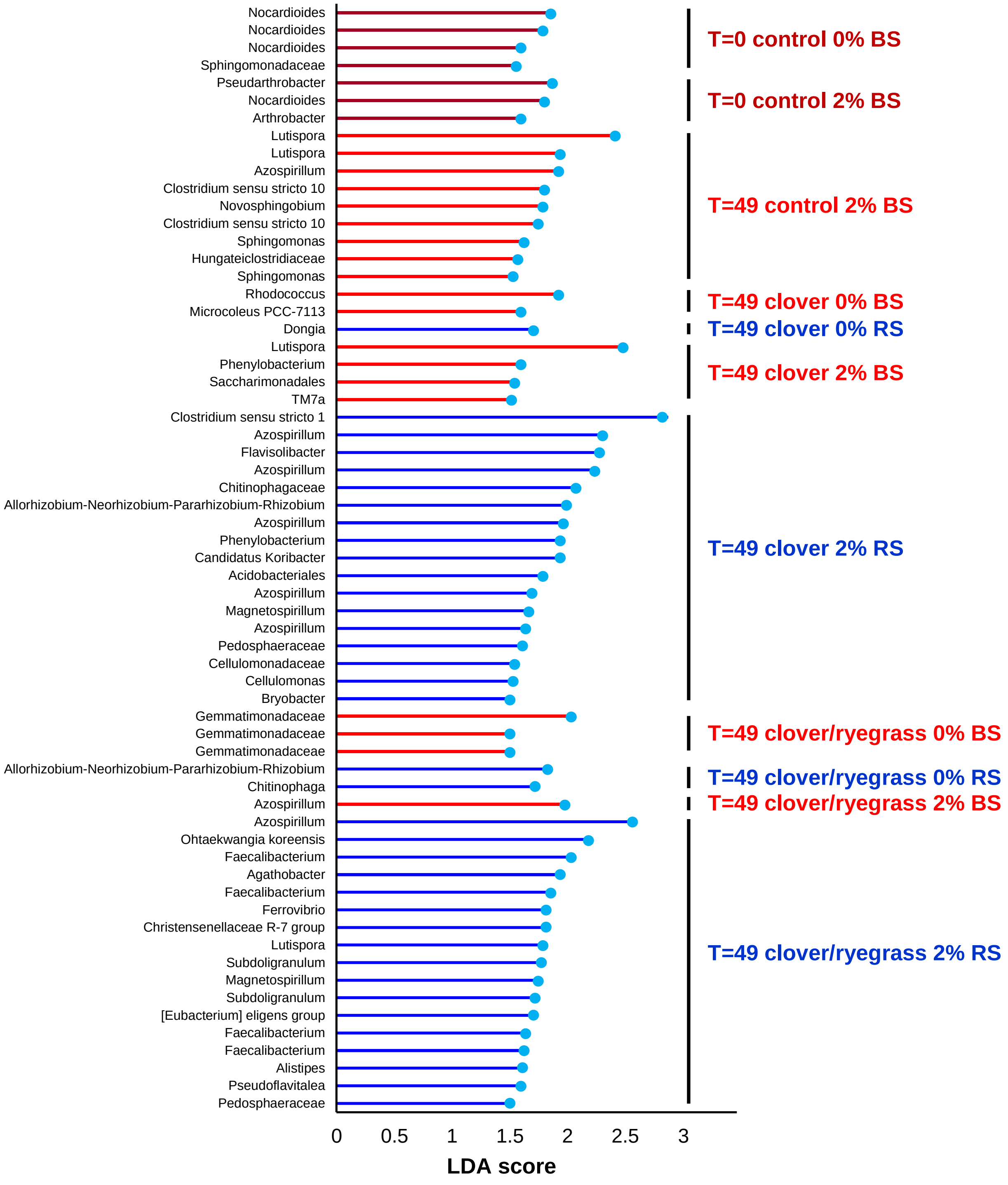


**Fig. S5** Biomarker ASVs identified by LefSe (>1.5% LDA) for each condition and time. BS is bulk soil, RS is rhizosphere soil, % diesel.

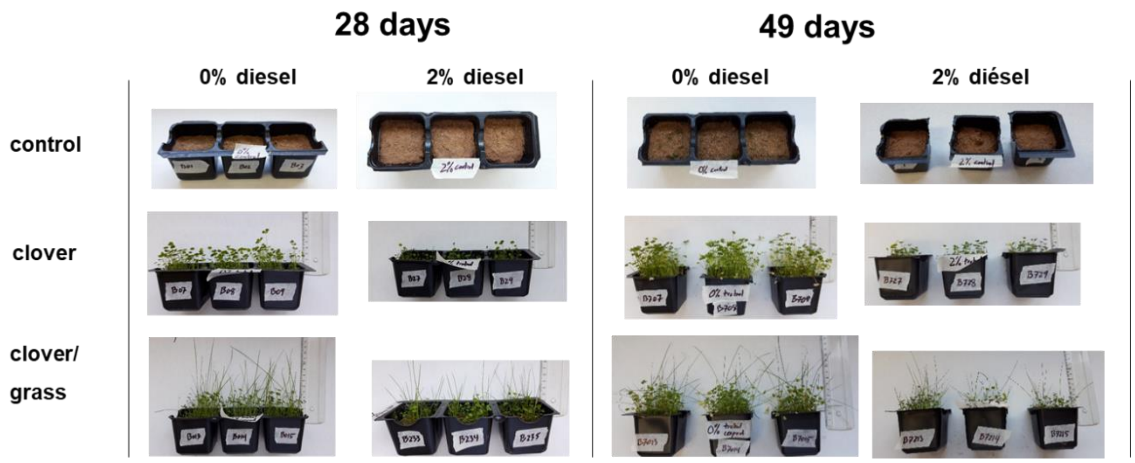


**Fig. S6** Aspect of clover and grass after 28 and 49 days in soil with or without 2% diesel. Rulers (0 cm) are aligned with the soil surface.
